# WDFY2 restrains matrix metalloprotease secretion and cell invasion by retention of VAMP3 in endosomal tubules

**DOI:** 10.1101/299610

**Authors:** Marte Sneeggen, Nina Marie Pedersen, Coen Campsteijn, Ellen Margrethe Haugsten, Harald Stenmark, Kay Oliver Schink

## Abstract

The endosomal FYVE- and WD40-domain-containing protein WDFY2 has been assigned a function as tumour suppressor, but its functional mechanism has remained elusive. Here we have used confocal, widefield and super-resolution fluorescence microscopy to show that WDFY2 localizes to the base of retromer-containing endosomal tubules by a mechanism that involves recognition of a specific pool of phosphatidylinositol 3-phosphate (PtdIns3P) by the WDFY2 FYVE domain. Affinity purification and mass spectrometry identified the v-SNARE VAMP3 as an interaction partner of WDFY2, and cellular knockout of WDFY2 caused a strong redistribution of VAMP3 into small vesicles near the plasma membrane. This was accompanied by increased secretion of the matrix metalloprotease MT1-MMP, enhanced degradation of extracellular matrix, and increased cell invasion. WDFY2 is frequently lost in metastatic cancers, most predominantly in ovarian and prostate cancer. We propose that WDFY2 acts as a tumor suppressor by serving as a gatekeeper for VAMP3 recycling.

## Introduction

One of the most life-threatening aspects of cancer is the ability of transformed cells to invade into the extracellular matrix (ECM) and neighbouring tissue to form metastases^1, 2^. Metastasis is often correlated with aggressive tumors and poor prognosis for the patient and therefore one of the leading causes of death by cancer^1^. Matrix metalloproteases (MMPs) play a critical role in progression of cancer by degrading and remodeling extracellular matrix, making the cells able to overcome tissue barriers, travel within the circulatory system before extravasating to produce a secondary tumor^3, 4^. MMPs such as MT1-MMP are internalized by clathrin-dependent and caveolar endocytosis^5^. After internalization, MT1-MMP is sorted in endosomal compartments, and a fraction is recycled back to the plasma membrane. How MMPs are sorted in endosomes is largely unknown. Sorting of endocytic cargos occurs at specialized tubular domains of early endosomes, and it is likely that sorting of MMPs occurs by similar mechanisms, however, the molecular factors which regulate this process are largely unknown.

WDFY2 has been described to reside on endocytic vesicles close to the plasma membrane^6^. It contains a lipid-binding FYVE domain and seven WD 40 repeats which can form a β-propeller. Potentially β-propellers can act as platforms for protein-protein interactions, but only few interactors of WDFY2 have been identified to date, and its cellular functions still remain to be elucidated ^7, 8^.

Here, we show that WDFY2 regulates exocytosis of MT1-MMP by controlling endosomal sorting of the v-SNARE VAMP3. WDFY2 localizes to Rab4-positive sorting endosomes and actin-stabilized endosome tubules. Here it interacts with VAMP3, which directs secretion of endosome-derived cargos, including MT1-MMP. We show that loss of WDFY2 leads to enhanced secretion of MT1-MMP and allows cells to actively invade into extracellular matrix. WDFY2 is frequently lost in metastatic tumours, but the link between this pathology and the molecular function of WDFY2 has not been elucidated.

## Results

### WDFY2 localizes to tubular regions of early endosomes

During endocytic trafficking, newly-formed vesicles undergo a maturation process. By changing the protein composition of their limiting membrane, they form distinct vesicle pools with specialized biochemical properties and functions. After internalization and uncoating, endocytic vesicles gain the early endocytic marker APPL1. APPL1 vesicles can mature into WDFY2 positive vesicles, and these vesicles can then further mature into early endosomes containing the canonical marker EEA1^9^. To date, the function of WDFY2 in the endocytic pathway is poorly characterized. To define the localization of WDFY2 in the endocytic pathway, we transiently transfected GFP-WDFY2 in RPE1 cells and performed structured illumination imaging (SIM) together with APPL1 and EEA1 visualized with antibodies (Figure 1a). Whereas APPL1-positive vesicles localize close to the plasma membrane^9^, we observed that WDFY2 localized to a pool of vesicles that was further from the plasma membrane and negative for APPL1 and EEA1. Towards the center of the cell, WDFY2 localized to EEA1 positive endosomes, however, it resided on distinct, EEA1-negative subdomains (Figure 1a).

**Figure 1:**
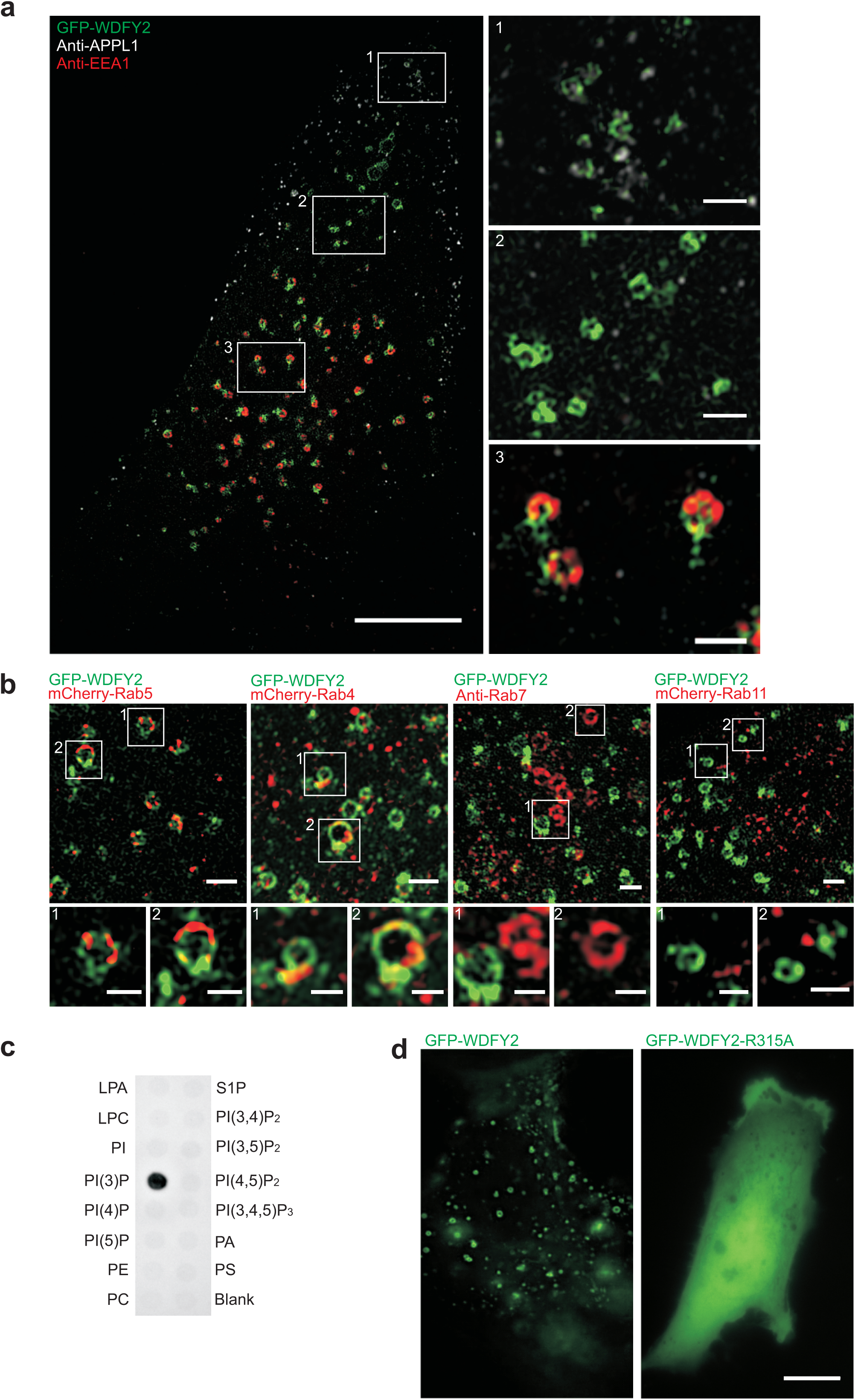
WDFY2 localization in the endocytic pathway. a. SIM image showing WDFY2 localization in relation to different markers of the early endocytic pathway. APPL1 (grey) was used as a marker for early vesicles and EEA1 (red) marks early endosome. GFP-WDFY2 (green) localizes to an independent vesicle pool (inset 2) and to subdomains on EEAl-labelled endosomes (inset 3). There is only limited overlap with APPL1 endosomes (inset 1). Scale bar: 10 μm, scale bar of insets 1 μm (n = 14 cells).
b. SIM images showing the localization of GFP-WDFY2 in relation to mCherry-Rab5 (n = 12 cells), mCherry-Rab4 (n = 13 cells), Anti-Rab7 (n = 14 cells) or mCherry-Rab11 (n = 14 cells). Scale bar: 1 μm, scale bar of insets: 0.5 μm.
c. Protein-Lipid overlay assay using purified full length WDFY2. WDFY2 binds with high selectivity to PtdIns(3)P.
d. Deconvolved widefield image showing GFP-WDFY2 localization to endosomes. GFP-WDFY2-R315A, a mutation in the binding site for PtdIns(3)P, abolishes the localization to endosomes and the protein is cytosolic. Scale bar: 10 μm (n=10 cells).

Earlier reports have proposed that WDFY2 does not localize to endosomes positive for the early endosomal marker Rab5 and therefore it has been suggested that WDFY2 marks a different set of endosomes which is distinct from those enriched in EEA^16, 9^. We therefore asked if WDFY2 localized together with any of the well-characterized Rab GTPases in the early endocytic pathway. SIM imaging was performed using a stable cell line expressing low levels of GFP-WDFY2 which was transiently transfected with mCherry-Rab4, -Rab5 and -Rab11. Endogenous Rab7 was visualized by antibody staining (Figure 1b). This imaging showed that WDFY2, in contrast to previous reports, localized to endosomes positive for Rab5, but resided on distinct, Rab5-negative subdomains of the endosomes (Figure 1b). We also observed that WDFY2 localized to endosomal regions positive for the fast-recycling marker Rab4. In contrast, we observed less overlap with the late-endosomal marker Rab7, and we could not observe co-localization with Rab11, a marker for the slow recycling pathway (Figure 1b).

Whereas most FYVE domains bind phosphatidylinositol-3-phosphate (PtdIns3P), also β-propellers have been described to be potential phosphoinositide (PI) binders^6^. We therefore asked if full-length WDFY2 was able to bind other PIs than PtdIns3P. By purifying full length WDFY2 and performing a protein lipid overlay assay we found that WDFY2 binds only to PtdIns3P with high specificity (Figure 1c). To test if this specificity is governed by the FYVE domain, we generated a stable cell line expressing GFP-WDFY2 with a point mutation in the conserved binding site for PtdIns3P (R315A), which abolishes PtdIns3P binding without distorting the overall FYVE structure^10^. This mutation completely disrupted the localization of WDFY2 to endosomes and led to a purely cytosolic localization (Figure 1d). Thus, we conclude that WDFY2 is recruited to endosomes by FYVE-dependent PtdIns3P binding.

Next we performed live-cell microscopy using the stable cell line expressing GFP-WDFY2 to trace the localization of WDFY2 in the endocytic pathway. Time-lapse movies showed GFP-WDFY2 localizing to endosomes, with a prominent localization to tubulating regions. These tubules emerged from WDFY2-rich domains on the endosomal membrane and showed strong accumulation of WDFY2 at their base (Figure 2a, Movie S1).

**Figure 2:**
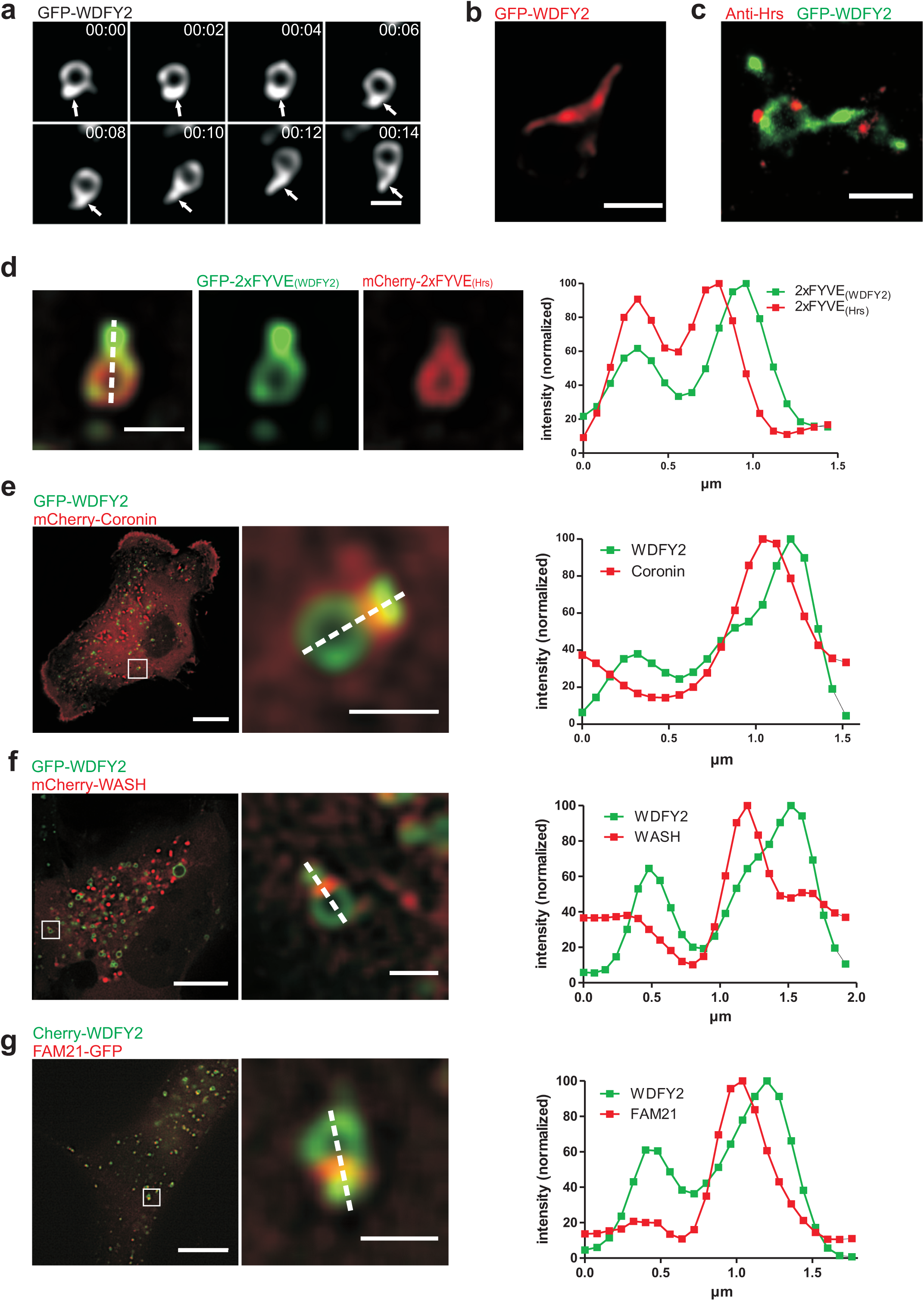
WDFY2 localizes to endosomal tubules a. Sequential images showing WDFY2 localization to newly formed tubular structures and WDFY2 accumulation on the base of the tubules. Shown are frames from a time-lapse sequence with images acquired every 2 seconds. Scale bar: 0.5 μm (n=200 endosomes)
b. STORM image showing WDFY2 localization to tubules, Scale bar: 0.5 μm Example image from n= 5 cells).
c. DNA-PAINT image showing GFP-WDFY2 (green) and Hrs (red) localization to endosomes. Scale bar: 0.5 μm Example image from n= 3 cells).
d. Distribution of GFP-2xFYVE_(WDFY2)_ and mCherry-2xFYVE_(Hrs)_ on a tubulating endosome. Scale bar: 1μm. A WDFY2-derived 2xFYVE probe shows a preference for tubulating membranes. The graph shows the normalized fluorescence intensity of 2xFYVE_(WDFY2)_ and mCherry-2xFYVE_(Hrs)_ along the indicated line (shown are representative data from n= 50 observed endosomes).
e. Localization of GFP-WDFY2 and mCherry-Coronin on a tubulating endosome. Scale bar 10 μm/1μm (inset). Coronin localizes to the base of WDFY2-positive tubules. The graph shows the normalized fluorescence intensity of GFP-WDFY2 and mCherry-Coronin along the indicated line (shown are representative data from n= 50 observed endosomes).
f. Localization of GFP-WDFY2 and mCherry-WASH on a tubulating endosome. Scale bar 10 μm/1μm (inset). WASH localizes to the base of WDFY2-positive tubules. The graph shows the normalized fluorescence intensity of GFP-WDFY2 and mCherry-WASH along the indicated line (shown are representative data from n= 50 observed endosomes).
g. Localization of mCherry-WDFY2 and GFP-Fam21 on a tubulating endosome. Scale bar 10 μm/1μm (inset). Fam21 localizes to the base of WDFY2-positive tubules. The graph shows the normalized fluorescence intensity of mCherry-WDFY2 and GFP-Fam21 along the indicated line (shown are representative data from n= 50 observed endosomes).

In order to gain super-resolved images of the tubular structures, we performed stochastic optical reconstruction microscopy (STORM) on fixed cells stably expressing GFP-WDFY2. This imaging showed an accumulation of WDFY2 at tubules emerging from the otherwise round endosome, as well as accumulations of WDFY2 at the base of these tubules. In contrast, the remaining limiting membrane of the endosomes showed only weak WDFY2 staining (Figure 2b). In addition, we performed DNA points accumulation for imaging in nanoscale topography (DNA-PAINT) microscopy of GFP-WDFY2 together with Hrs, a protein involved in protein sorting on early endosomes^11, 12^. WDFY2 labels both the limiting membrane and endosomal tubules, whereas Hrs localized only to microdomains of the vesicles and could not be detected on tubules positive for WDFY2 (Figure 2c).

We next asked if this tubular localization is based on the PtdIns3P-binding FYVE domain or other features of WDFY2. To this end, we generated a tandem PtdIns3P-binding probe based on the FYVE domain of WDFY2 (2xFYVE) and compared it to the widely used Hrs-derived 2xFYVE probe^13^. Surprisingly, whereas Hrs 2xFYVE localized only weakly to tubular structures, the WDFY2 derived 2xFYVE probe showed a strong preference for tubular structures (Figure 2d). This suggests that the localization of WDFY2 to small vesicles and endosomal tubules is controlled by its FYVE domain. Moreover, it highlights that different FYVE domains, despite binding to the same PI, can show preferences for distinct lipid subpopulations, based on physical or biochemical properties of the membrane.

Localization of WDFY2 to endosomal tubules has not been reported before and we therefore proceeded to characterize these structures in more detail. Endosomes give rise to different classes of tubules which can mediate sorting and transport of cargos. Previous reports have shown that actin-dependent, relatively stable tubules are involved in the sorting of slowly diffusing cargos such as β2-adrenergic receptors, whereas bulk recycling of freely diffusing cargo such as transferrin receptors (TfRs) happens via short-lived tubules^14^. We therefore asked to which tubule class WDFY2 localizes. We found that GFP-WDFY2-labelled tubules were relatively long-lived and showed accumulations of the actin organizing proteins Coronin and WASH at their base (Figure 2e,f, Movie S2)^15, 16^. We also found that the WASH-interacting protein FAM21 and subunits of the retromer cargo recognition complex - VPS26 and VPS35 (Figure 2g and Figure S1a,b)^16, 17^, localized to the base of GFP-WDFY2-labelled tubules. Depolymerization of actin with Latrunculin caused rapid changes in the morphology of GFP-WDFY2-labelled tubules. We observed that, immediately after addition of Latrunculin, endosomal tubules started to hyper-tubulate and pinched off as large fragments (Movie S3). After this initial reaction, tubulation ceased, in line with earlier reports^14^. This suggests that WDFY2 localizes to actin-stabilized tubules which are involved in specific sorting of cargo. Together with the colocalization with Rab4, this suggests that WDFY2 could be involved in controlling recycling of endocytosed cargos.

### WDFY2 interacts with the v-SNARE VAMP3

Little is known about the function of WDFY2 in the endocytic pathway, and few WDFY2 interaction partners are described so far^7, 8^. To further elucidate the function of WDFY2 we therefore set out to identify putative interaction partners. GFP-Trap immuno-precipitates from cell lines stably expressing GFP-WDFY2 or GFP as a control were analyzed by quantitative mass spectrometry (Supplemental Table S3). We decided to focus on proteins that were at least 50-fold enriched in GFP-WDFY2 purifications and searched for candidates that could help understand the function of WDFY2 in the endocytic system (Figure 3a).

**Figure3:**
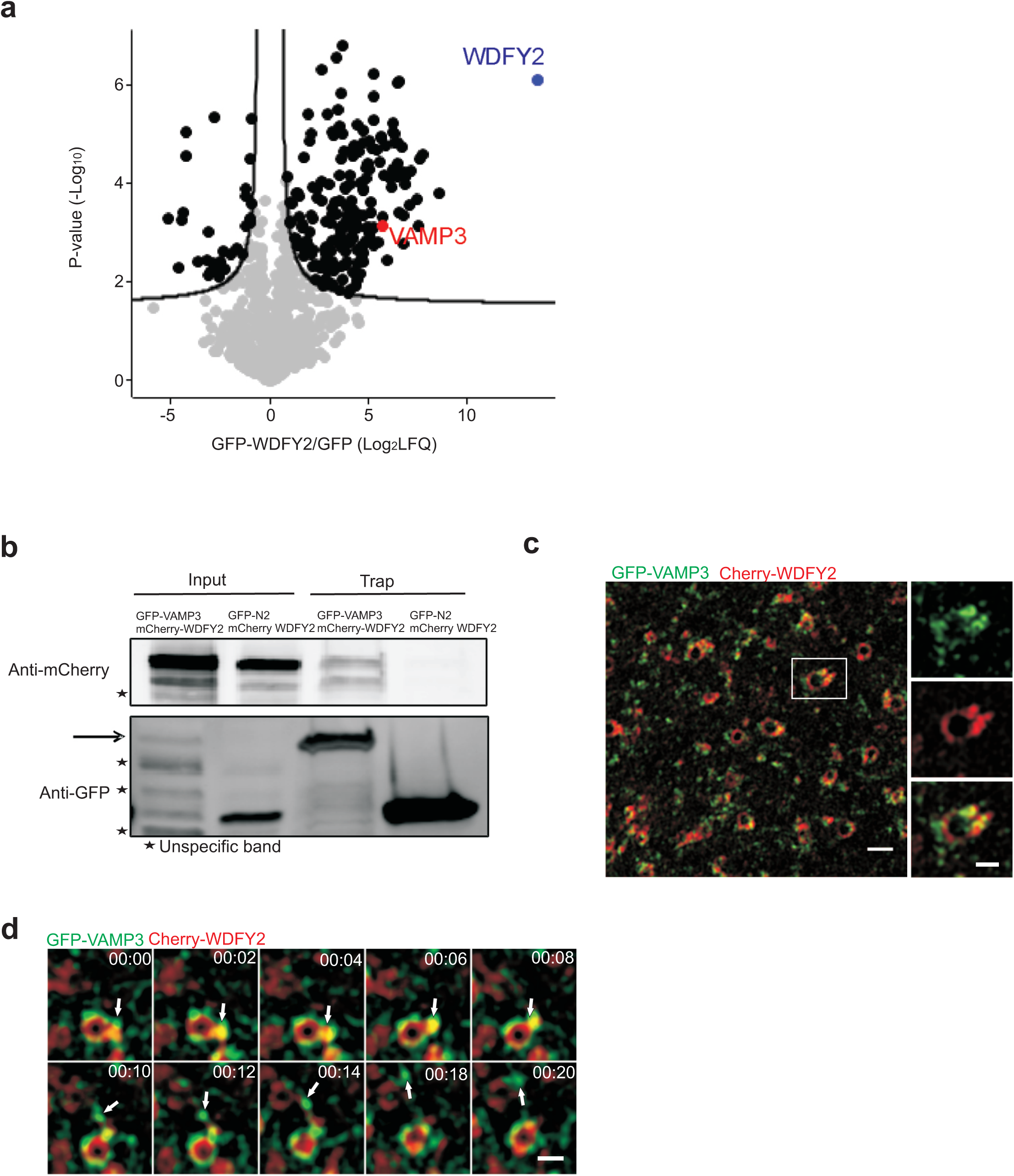
WDFY2 interacts and colocalizes with VAMP3 a. Volcano plot identifying enriched proteins isolated from cells expressing GFP-WDFY2 by GFP-trap affinity purification of cell lysates in comparison to GFP expressing control cells. Labelfree quantification (LFQ) ratios of GFP-WDFY2/GFP pulldowns were plotted against the p-value, with the permutation-based FDR threshold (0.01) indicated by the black lines.
b. GFP affinity purification of GFP-VAMP3 and mCherry-WDFY2. Affinity purification of GFP-VAMP3 can co-precipitate mCherry-WDFY2, whereas isolation of GFP alone does not coprecipitate mCherry-WDFY2 (n= 3 experiments).
c. SIM image showing GFP-VAMP3 and Cherry-WDFY2 localization to endosomes and endosomal tubules. Scale bar overview image: 1 μm, scale bar of inset image: 0.5 μm (n=13 cells).
d. Sequential images showing GFP-VAMP3 and Cherry-WDFY2 localization to endosomes and endosomal tubules. Shown are frames from a time-lapse movie with images acquired every 2 seconds. Scale bar: 0.5 μm (n= 10 cells).

Using this method, we identified the v-SNARE protein VAMP3 as a potential interaction partner, which showed 52-fold enrichment in the mass spectrometry analysis (Figure 3a). VAMP3 has been shown to be present on recycling endosomes and involved in Transferrin recycling^18^. It can also bind to the plasma membrane through the t-SNAREs syntaxin1, syntaxin4, SNAP23 and SNAP25^19^. VAMP3 is also usually segregated into tubular membranes where it facilitates fusion with the endocytic compartment and the Golgi apparatus^18^. To verify this interaction, we performed GFP affinity purifications followed by western blotting. This experiment confirmed the interaction of WDFY2 and VAMP3 (Figure 3b).

Next, we asked whether WDFY2 and VAMP3 colocalized on the same vesicles and if VAMP3 resides on WDFY2 positive tubules. To this end, we performed SIM imaging of cells transiently transfected with GFP-VAMP3 and Cherry-WDFY2 (Figure 3c). This revealed that WDFY2 and VAMP3 were localized to the same endosomes. In addition, VAMP3 could also be found on WDFY2 negative vesicles (Figure 3c). Live cell imaging showed that VAMP3 localizes to WDFY2 positive tubules (Figure 3d). We observed that from these tubules, vesicles positive for VAMP3 and WDFY2 were pinched off. Directly after pinching off, WDFY2 left the newly-formed vesicles, whereas VAMP3 remained.

### VAMP3 is redistributed to the leading edge of WDFY2 knockout cells

After VAMP3 vesicles are pinched off from WDFY2-containing tubules, they are presumably transported to different compartments where VAMP3 can mediate membrane fusion. In wild-type RPE1 cells, VAMP3 preferentially localized to vesicles which were dispersed throughout the cell and showed an accumulation of vesicles in the area surrounding the Golgi apparatus (Figure 4a).

**Figure 4:**
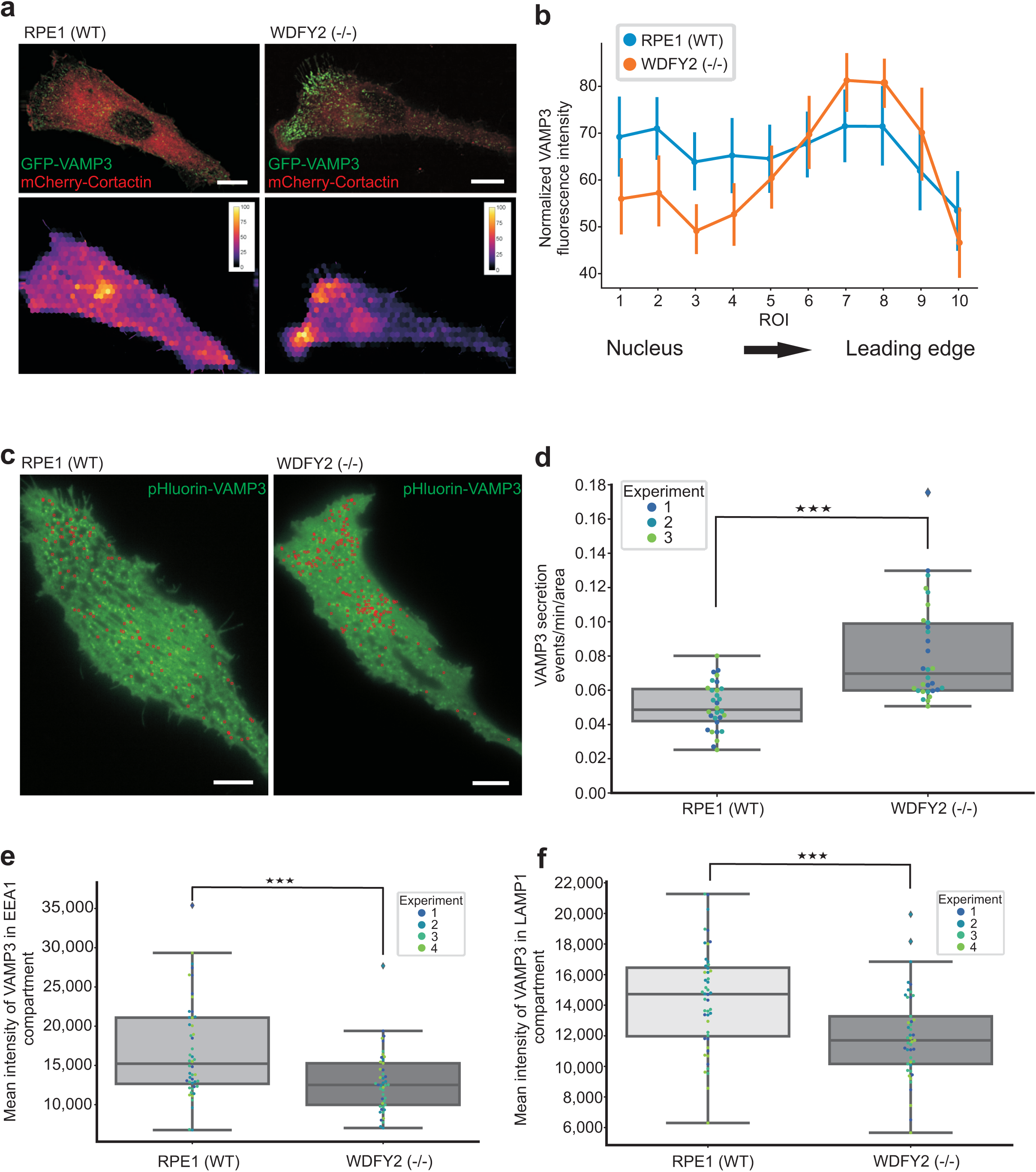
WDFY2 controls intracellular VAMP3 distribution a. Deconvolved widefield images of RPE and WDFY2 (-/-) cells transiently expressing GFP-VAMP3 and Cherry-Cortactin. Scale bar: 10 μm (n= 30 cells per condition). Hexagonal superpixel image showing VAMP3 distribution in the cells shown above. Mean intensities of hexagonal ROIs were extracted and the ROI filled with the corresponding mean value, thereby generating superpixels.
b. Distribution of VAMP3 from the nucleus to the leading edge. Cells were transfected with VAMP3 and stained for Cortactin to identify leading edges. A line ROI from the nucleus to the leading edge was drawn, and evenly spaced ROIs were automatically generated. Mean intensity of VAMP3 in each ROI was extracted, normalized for each cell, and then plotted, (n=30 cells for each condition) shown are mean and 95% CI.
c. TIRF micrograph of RPE1 (WT) and WDFY2 (-/-) cells transfected with VAMP3-pHluorin. Individual secretion events (summed over a two minute interval) are indicated with a red circle. Scale bar: 10 μm (n=30 cells per condition).
d. Quantification of VAMP3-pHluorin secretion in RPE1 (WT) and WDFY2 (-/-) cells. Shown are events from 3 experiments (10 cells per experiment per condition). Secretion events per minute are shown normalized to cell area. Student’s unpaired t-test, n=30, p=0.0000029.
e. Quantification of GFP-VAMP3 intensity in EEA1-positive endosomes in RPE1 (WT) and WDFY2 (-/-) cells. Plotted is the mean intensity within EEA1-positive endosomes per cell from 4 experiments (10,12,15,10 cells for (RPE1 (WT)), 10,10,15,10 cells for (WDFY2 (-/-)). Student’s unpaired t-test, n=47/45, p= 0.00035.
f. Quantification of GFP-VAMP3 intensity in Lamp1-positive endosomes in RPE1 (WT) and WDFY2 (-/-) cells. Plotted is the mean intensity within Lamp1-positive vesicles per cell from 4 experiments (10,12,15,10 cells for (RPE1 (WT), 10,10,15,10 cells for (WDFY2 (-/-)). Student’s unpaired t-test, n=47/45, p= 0.00026.

We therefore asked whether VAMP3 localization was affected by the absence of WDFY2. To this end, we generated WDFY2 knockout RPE1 cells using CRISPR/Cas9 (Figure S2)^20^.

In order to study the effect of WDFY2 on the distribution of VAMP3, RPE1 wildtype (WT) cells and the RPE1 WDFY2 (-/-) cells were transfected with GFP-VAMP3. We observed that in knockout cells, GFP-VAMP3 vesicles clustered just beneath the leading edge of migrating cells, which was labelled with mCherry-Cortactin (Figure 4a). In RPE1 (WT) cells, this localization was not as prominent. In order to quantify this localization, we generated a set of 10 regions of interest (ROIs) from the nucleus to the leading edge, each encompassing 10 % of the distance. The mean VAMP3 intensity of each ROI was extracted and plotted (Figure S3a). These measurements showed an even distribution of VAMP3 in WT cells, with small peaks close to the nucleus and the leading edge (Figure 4b). In comparison, WDFY2 (-/-) cells showed a dramatic redistribution of VAMP3, with low concentrations of VAMP3 localized close to the nucleus and a strong accumulation of VAMP3 close to the leading edge of the cell.

### WDFY2 controls VAMP3-dependent secretion

We then asked if this accumulation of vesicles at the leading edge could result in higher rates of VAMP3-driven endocytic recycling to the plasma membrane. To this end, we utilized a pHluorin-based exocytosis assay^21^. We transfected RPE1 (WT) and WDFY2 (-/-) cells with VAMP3-pHluorin. In cells expressing this construct, the pH sensitive pHluorin tag is facing the lumen of endocytic and secretory vesicles and will be exposed at the cell surface as the vesicle fuses with the plasma membrane. In the acidic environments of endosomes the pHluorin fluorophore will be quenched. Once the recycling vesicle has fused with the plasma membrane the fluorophore will be unquenched. Cells were imaged every second for two minutes using total internal reflection fluorescence imaging (TIRF) and bright flashes, indicating fusion events within the plasma membrane, were counted manually using Fiji. We observed that the recycling rate was significantly elevated in the WDFY2 (-/-) cells compared to the RPE1 (WT) cells, indicating that loss of WDFY2 does not only lead to redistribution of VAMP3 vesicles to the leading edge but also results in higher recycling rates. This is presumably the result of more VAMP3-containing recycling vesicles at the leading edge (Figure 4 c,d, Movie S4).

VAMP3 is recycled via endosomal compartments, and based on the localization of WDFY2 to tubules, an attractive hypothesis was that lack of WDFY2 could lead to changes in VAMP3 recycling. To test this hypothesis, we transfected RPE1 (WT) and WDFY2 (-/-) cells with GFP-VAMP3 and measured the distribution with different endocytic compartments. We observed that EEA1-positive early endosomes in WDFY2 (-/-) cells showed reduced levels of VAMP3 (Figure 4e), suggesting that either less VAMP3 arrives at the early endosomes or that recycling occurs faster. LAMP1-labelled vesicles also showed reduced VAMP3 levels (Figure 4f), indicating that the observed depletion of VAMP3 in early endosomes was not a result of a faster transport of cargo towards the degradative pathway.

### Loss of WDFY2 leads to enhanced secretion of matrix metalloproteases

In order to characterize the cellular function of WDFY2, we set out to find a cargo transported of the vesicles that were accumulating at the leading edge in WDFY2 (-/-) cells. VAMP3-positive vesicles have been shown to transport the membrane anchored matrix metalloprotease MT1-MMP to the plasma membrane^4, 22^. MT1-MMP has been reported to be transported via Rab4-dependent fast endocytic recycling, and Rab5A can redirect MT1-MMP to invadosomes in order to allow ECM degradation and cell invasion^23^.

To investigate whether MT1-MMP could be a potential cargo for the WDFY2 positive tubules and VAMP3 containing vesicles, we performed live-cell imaging of RPE1 cells transfected with mCherry-WDFY2 or VAMP3 in combination with GFP-MT1-MMP (Figure 5a, b, Movie S5). MT1-MMP clearly colocalized with both WDFY2 and VAMP3. We also observed that MT1-MMP is sorted into WDFY2-labelled tubules and transported in VAMP3 positive vesicles (Figure 5b).

**Figure 5:**
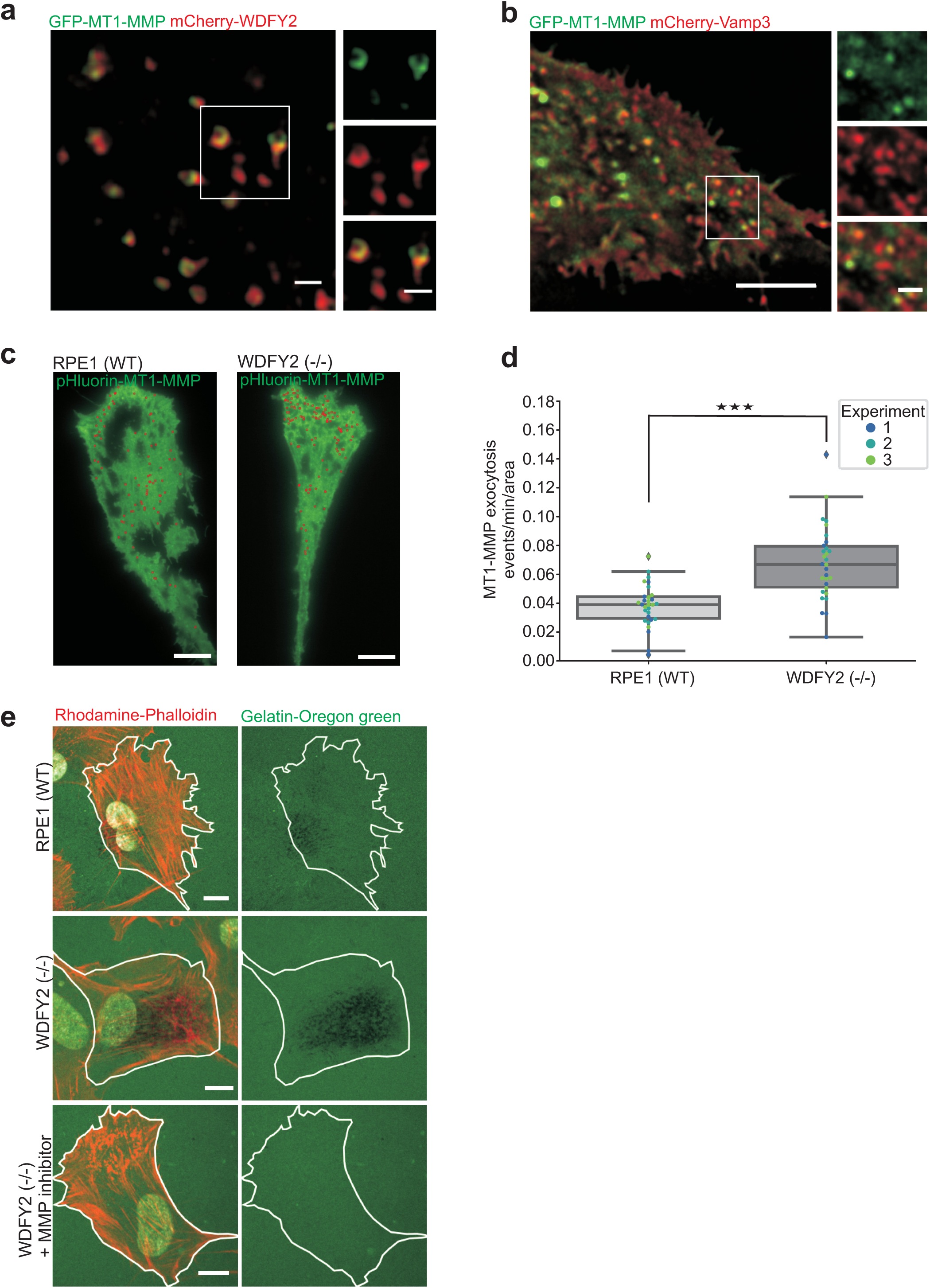
WDFY2 controls MT1-MMP trafficking and extracellular matrix degradation a. Deconvolved widefield image showing localization of GFP-MT1-MMP to mCherry-WDFY2-positive endosomes and endosomes tubules. Scale bar: 1 μm, scale bar of inset: 1 μm.
b. Deconvolved widefield image showing GFP-MT1-MMP and mCherry-VAMP3 colocalization at vesicles. Scale bar: 5 μm, scale bar of inset: 1 μm.
c. TIRF micrograph of RPE1 (WT) and WDFY2 (-/-) cells transfected with pHluorin-MT1-MMP. Individual exocytic events (summed over two minutes) are indicated with a red circle. Scale bar: 10 μm (n=30 cells per condition).
d. Quantification of pHluorin-MT1-MMP exocytosis in RPE1 (WT) and WDFY2 (-/-) cells. Shown are events from 3 experiments (10 cells per experiments for per condition). Exocytosis events per minute are shown normalized to cell area. Student’s unpaired t-test,n=30, p=0.000001.
e. Confocal micrographs showing degradation of a fluorescent gelatin layer, indicated by dark areas, by RPE1 (WT), WDFY2 (-/-) and MT1-MMP-inhibitor treated WDFY2 (-/-) cells.

Based on these observations, we asked if the secretion rate of MT1-MMP is affected by the loss of WDFY2. To investigate this, we performed TIRF live-cell microscopy of RPE1 (WT) and WDFY2 (-/-) cells expressing pHluorin-MT1-MMP^24^. We observed that the rate of MT1-MMP exocytosis was strongly increased in the absence of WDFY2 (Figure 5c,d, Movie S6). The exocytosis rate and the observed change in WDFY2 (-/-) cells showed the same tendency as observed in the VAMP3 exocytosis experiment, further supporting the notion that VAMP3-driven exocytosis of MT1-MMP is controlled by WDFY2.

Based on our observation that more MT1-MMP was secreted from cells lacking WDFY2, we asked if these cells also were able to degrade more ECM. To this end, we assayed degradation of gelatin labelled with the fluorescent dye Oregon green. RPE1 (WT) and WDFY2 (-/-) cells were seeded on coverslips coated with a layer of fluorescent gelatin^25^. Upon six hours at 37°C the cells were fixed and stained with the Actin marker Phalloidin to visualize the cells whereas gelatin degradation was visualized as black areas in the gelatin layer. By confocal imaging we observed that WDFY2 (-/-) cells degraded visibly more gelatin in comparison to RPE1 (WT) cells (Figure 5e). In both cell lines, incubation with the MMP inhibitor GM6001 resulted in a complete lack of degradation, indicating that the observed effects were indeed based on secretion and function of MT1-MMP (Figure 5e, data not shown for RPE (WT)). This suggests that loss of WDFY2 does not only enhance exocytosis of pHluorin-MT1-MMP, but also regulates trafficking of MT1-MMP and thereby its ability to degrade ECM.

### Loss of WDFY2 causes increased cell invasion

Cancer cells often show elevated levels of MMPs and increased secretion, which allows them to invade neighbouring tissues and establish metastases^5^. The elevated MT1-MMP secretion and degradation of ECM in WDFY2 (-/-) cells led us to ask if this allowed cells lacking WDFY2 to invade into 3D matrices. To this end, we performed 3D inverted invasion assays in Matrigel^26^. RPE1 (WT) and WDFY2 (-/-) cells were seeded on to the bottom of a Transwell filter and allowed to migrate through the filter and into Matrigel supplemented with fibronectin along a gradient of serum and hepatocyte growth factor (HGF). Invading cells were stained with Calcein AM and monitored by confocal microscopy. Invasion was measured according to Oppelt *et.al*, 2014^26^. We observed that WDFY2 (-/-) cells were able to invade further into Matrigel in comparison to wild type control cells (Figure 6a,b,c). This observation was supported by siRNA-based depletion of WDFY2, which showed the same phenotype as a full knockout (Figure S4). Importantly, the effect on WDFY2 depletion could be rescued by stable expression of a siRNA-resistant WDFY2 construct, confirming the involvement of WDFY2 in cell invasion.

**Figure 6:**
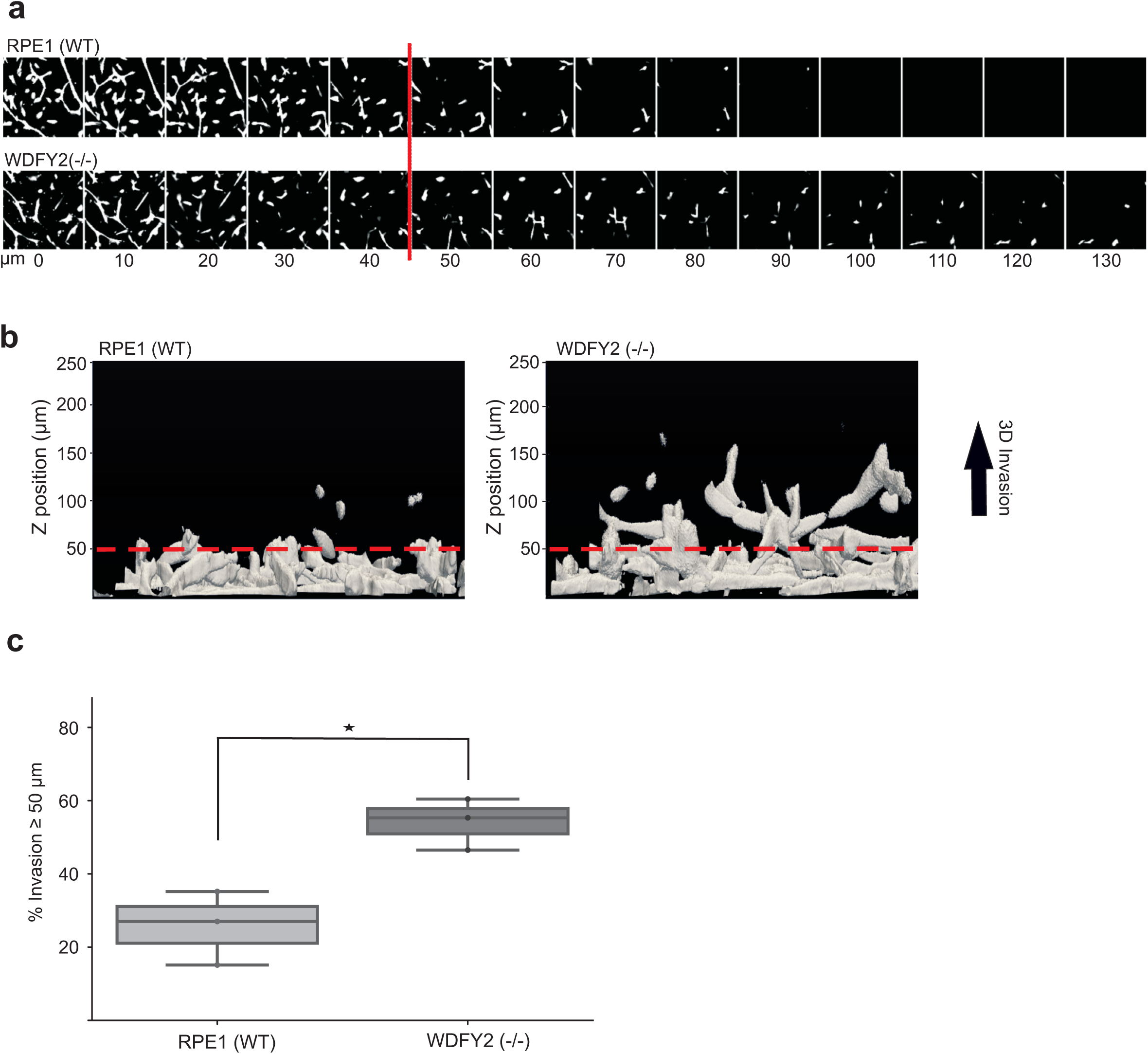
WDFY2 controls cell migration and invasion. a. Optical sections (Δz=10 μm) of RPE (WT) and WDFY2 (-/-) cells stained with Calcein-AM invading into fibronectin-supplemented Matrigel^™^. The red line indicates the z-axis threshold (50 μm) defining invading cells.
b. Orthogonal view of a 3D reconstruction of RPE (WT) and WDFY2(-/-) invading into fibronectin-supplemented Matrigel^™^. The red line indicates the z-axis threshold (50 μm) defining invading cells.
c. Quantification of invasion of RPE1 (WT) and WDFY2 (-/-) cells into fibronectin-supplemented Matrigel^™^. Invasion was quantified as described in the experimental section. Plotted data points indicate measurements derived from individual confocal z-stacks, color-coded per experiment (15 z-stacks per experiment, n= 3 experiments). Student’s t-test, p=0.016.

## Discussion

In this study, we provide evidence for a novel mode of action for a tumour suppressor, namely by restricting endocytic recycling of a MMP and thereby preventing cell invasion. We show that the endosomal protein WDFY2 regulates endocytic recycling to the plasma membrane of MT1-MMP via a VAMP3-dependent mechanism, and that knockout of WDFY2 causes increased degradation of extracellular matrix accompanied by increased cell invasion.

We found that WDFY2 labeled a subpopulation of endosomes distinct from APPL1 and EEA1-positive endosomes, which was positive for the recycling marker Rab4. It has previously been reported that WDFY2-carrying vesicles are Rab5-negative but are in close association with Rab5-positive endosomes ^6^. In contrast to these reports, we observed that both Rab5 and WDFY2 were present on the same vesicle, but resided in distinct subdomains, a feature that might have been non-resolvable in previous studies. One of the most striking features was that WDFY2 localized to tubules emanating from EEA1-positive early endosomes which are positive for the retromer complex and the actin bundling protein, Coronin. We observed that vesicles were formed from these tubules which carried recycling cargoes such as VAMP3 and MT1-MMP. Together with the colocalization with Rab4, this suggests that WDFY2 could localize to structures that regulate cargo sorting for recycling.

Localization of WDFY2 was dependent on a functional FYVE domain and the full-length protein binds with high specificity to the phosphoinositide PtdIns3P. We also observed a clear selectivity of the WDFY2-derived 2xFYVE probe for PtdIns3P (not shown). Surprisingly, when probing PtdIns3P localization with the canonical Hrs-derived 2xFYVE probe, we observed only poor labeling of tubular structures on endosomes, whereas a WDFY2-derived 2xFYVE probe showed a clear preference for tubular endosomes. This argues that the WDFY2 FYVE domain has different binding properties in comparison to the Hrs FYVE domain; while both can bind PtdIns3P, they detect distinct pools. This observation has broader implications for the targeting of proteins by lipid-binding FYVE domains. It demonstrates that localization of these proteins does not only depend on the pure presence of PtdIns3P, but that FYVE domains can recognize and preferentially bind to PtdIns3P in a specific context. Sensing of this context could depend on the local properties of the host membrane, for example local curvature, membrane fluidity or the presence of other lipid species, which could allow a differential exposure of the PtdIns3P head group. As FYVE domains bind to PtdIns3P-containing membranes by inserting a hydrophobic loop structure^27^, this mechanism could support the sensing of PtdIns3P in different environments. Our findings also demonstrate the limitations of the current generation of PtdIns3P sensors, and our study provides a new sensor able to detect PtdIns3P on tubular membrane structures.

We identified the v-SNARE VAMP3 as an interactor of WDFY2 and established that cells deleted for WDFY2 show enhanced accumulation of VAMP3-labelled vesicles in the peripheral regions of the cell. WDFY2 (-/-) cells also showed higher recycling rates of VAMP3-positive vesicles. A cargo of VAMP3 vesicles, MT1-MMP^22^, showed the same behavior, indicating that the overall trafficking of VAMP3positive vesicles is controlled by WDFY2. This raises the question how WDFY2 could influence VAMP3-dependent trafficking. WDFY2 and VAMP3 colocalized at tubular structures, suggesting that the observed interaction of VAMP3 and WDFY2 occurs at these membrane structures. Similarly, MT1-MMP is sorted via the endolysosomal pathway, however, less is known about its trafficking^5^.

Sorting and retrieval of endocytosed cargo is one of the fundamental functions of the endocytic system. Tubular membrane structures are thought to play a major role in this process; however, how their cargo is selected is largely unknown^14^. On early endosomes, several sorting pathways have been described which are involved in trafficking of different cargos. Some cargos, such as TfRs, are sorted into short-lived tubules, and it has been shown that TfR can be sorted into these structures by fast diffusion^14^. In contrast to this, slow-diffusing cargos like β2-adrenergic receptors are sorted into stable actin-dependent membrane tubules^14^. These tubules are relatively long-lived and can form vesicles which are recycled back to the plasma membrane. One of their key characteristics is that they recruit actin and actin-binding proteins such as Coronin and WASH which stabilize the tubules and allow them to form stable sorting platforms^14^. The tubular structures labeled by WDFY2 were positive for Coronin and WASH and sensitive to actin disruption by Latrunculin, suggesting that they could represent actin-stabilized tubules.

Loss of WDFY2 led to a peripheral localization and enhanced secretion of VAMP3-positive vesicles and their cargo, MT1-MMP. How could the loss of WDFY2 lead to these changes? Based on the characteristics of WDFY2-positive tubules, it is an attractive hypothesis that WDFY2 normally sorts VAMP3 into stable, actin-dependent recycling tubules and thereby restricts the number of v-SNARE molecules on recycling vesicles. It is also likely that a significant portion of VAMP3 would be retained in endosomes and thus finally ends up in lysosomes. A loss of WDFY2 would abolish this control and allow VAMP3 to be sorted into bulk recycling tubules, leading to increased recycling.

Consistent with this, WDFY2 (-/-) cells showed less VAMP3 in both EEA1 and LAMP1 vesicles, supporting the notion that loss of WDFY2 leads to altered recycling of VAMP3. The increased secretion of MT1-MMP in the absence of WDFY2 is likely a direct consequence of altered VAMP3 sorting by providing more recycling vesicles for fusion with the plasma membrane. It is not clear how the number of v-SNARE molecules could influence the rate of membrane fusion, but one explanation could be that more newly-formed vesicles gain VAMP3 and thereby a secretory identity. VAMP3 can also recruit the phosphoinositide 4-kinase PI4K2A^18^. PI4K2A has been shown to mediate PI conversion and generation of Phosphatidylinositol-4-phosphate (PtdIns4P) during the formation of recycling vesicles^28^. As PtdIns4P is involved in anterograde transport of secretory vesicles, VAMP3-based recruitment of PI4K2A and altered levels of PtdIns4P could also explain the altered distribution of VAMP3-positive vesicles in WDFY2 (-/-) cells, as higher VAMP3 levels would recruit more PI4K2A and thereby establish PtdIns4 and a secretory identity on these vesicles.

Several lines of evidence suggest that WDFY2 can act as a tumour suppressor. A screen of the CBioportal cancer genome database shows that WDFY2 is frequently (in up to 14% of cases) lost in metastatic cancers (Figure 7a)^29, 30^. An earlier study reported a fusion gene consisting of *CDKN2D* and *WDFY2*, which occurs frequently in high-grade serous ovarian cancer (in 20 % of all HG-SC tumors)^31^. The fusion leads to expression of a truncated WDFY2 protein^31^. It is likely that this fusion protein would be unable to control VAMP3 trafficking as part of the first WD repeat is missing and the truncated protein would not form a functional β-propeller. Overexpression of WDFY2 has been proposed to reduce cell migration in an AKT1-dependent manner in prostate cancer, however, a cell biological explanation for this observation is missing so far^32^.

**Figure 7:**
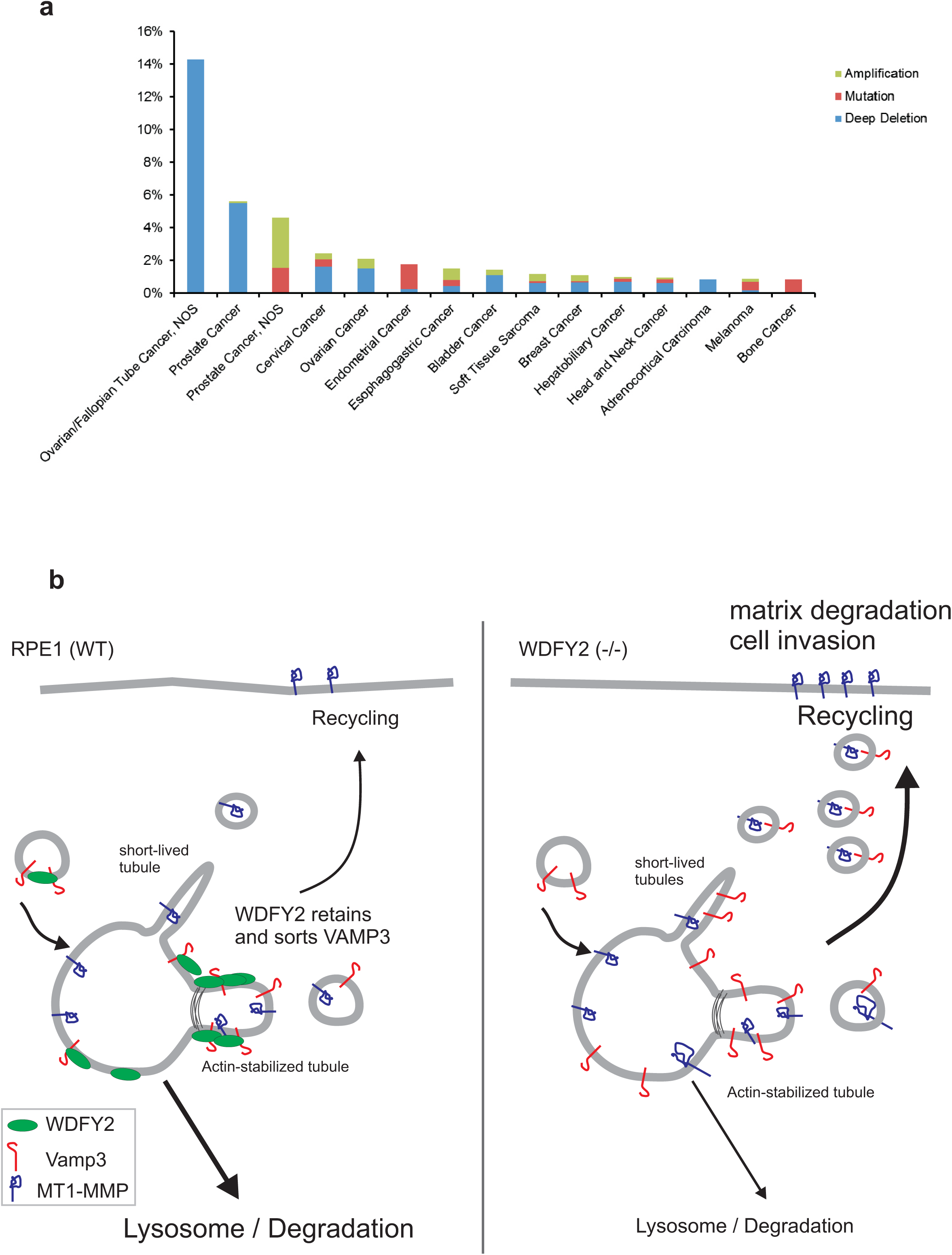
WDFY2 controls cancer cell invasion a. WDFY2 is frequently deep deleted in cancer samples. The graph shows data extracted from the cBioPortal for Cancer Genomics, alteration frequency is grouped by cancer type and plotted.
b. Model of WDFY2 action in the endocytic system. WDFY2 binds to the v-SNARE VAMP3 on endosomes and sorts it into actin-stabilized tubules. Binding to WDFY2 prevents recycling of VAMP3 by short-lived endosome tubules, and a major portion of VAMP3 is retained on endosomes and is trafficked to the lysosomal degradation pathway. Loss of WDFY2 allows enhanced recycling of VAMP3 and the generation of more vesicles with a secretory identity. These vesicles carry MT1-MMP. Elevated secretion of MT1-MMP allows enhanced ECM degradation and cell invasion.

Our data show that WDFY2 controls trafficking and secretion of the matrix metalloprotease MT1-MMP. We show that loss of WDFY2 leads to enhanced secretion of MT1-MMP, which allows cells to invade into 3D matrices. Correspondingly, a deletion of WDFY2 in cancer cells would enable them to migrate through the ECM and provide a higher metastatic potential, which correlates well with the finding that WDFY2 is frequently lost in metastatic cancers. Thus, WDFY2 could normally limit cell invasion by controlling VAMP3-dependent secretion, and a loss of this traffic controller would lead to significantly enhanced cell migration and metastatic potential. VAMP3 has been implicated as a major regulator of cell migration^33^. Migrating macrophages show a redistribution of VAMP3, similar to what we observed in WDFY2-depleted cells, and it has been suggested that this redistribution controls lamellipodia formation and cell migration^34^. Also in cancer cells, VAMP3 has been shown to be involved in the secretion of MT1-MMP^22^. We conclude that WDFY2 acts as a key traffic controller which curbs cell invasion by regulating VAMP3 trafficking and MT1-MMP secretion, and the loss of this safety mechanism.

## Material and Methods

### Antibodies

The following antibodies were used: Human anti EEA1 provided by Ban-Hock Toh (Monash University), Rabbit anti APPL1 D83H4 from cell signaling (3858S), Rabbit anti Rab7 was from cell signaling (9367), Mouse anti GFP was from Roche (11814 460001), RFP-booster atto 594 was from Chromotek (rba594), Rabbit antibody against Hrs have been described previously^35^, Rabbit anti LAMP1 was from Sigma-Aldrich (L1418), Rhodamine Phalloidin (R415) and Hoechst 33342 (H3570) was from Invitrogen molecular Probes, Goat anti Vps35 (ab10099), Rabbit anti Vps26 (ab23892) were from Abcam. Goat anti Cherry was from Acris Antibodies (AB0040-200). HRP-conjugated anti-GST antibody was from GE Healthcare (RPN1236). All secondary antibodies used for IF were obtained from Jacksons ImmunoResearch Laboratories and secondary antibodies used for western blotting were obtained from LI-COR Bioscience GmbH.

### Plasmids

pmCherry-Rab11a was a gift from Jim Norman^36^, pCDNA-pHlourin_MT-MMP was gift from Philippe Chavrier^37^, for some experiments, the pHluorin tag was exchanged with eGFP, Fam21-GFP (pEGFP-N1-3) was a gift from Dr. Matthew Seaman^16^. The following plasmids were obtained from Addgene: pmCherry-Rab4 (55125) and pmCherry-WASH1-C-18 (55162) were a gift from Michael Davidson, pEGFP-VAMP3 (42310) was a gift from Thierry Galli^38^. pmCherry-Cortactin (27676) and Coronin1B-mCherry (27694) were a gift from Christien Merrifield^39^. pX458 (48138) was a gift from Feng Zhang^20^. To create pVamp3-pHluorin, Vamp3 and superecliptic pHluorin were PCR-amplified and cloned into a pEGFP-N1-derived backbone.

All other plasmids were generated using standard cloning procedures and are listed in Supplemental Table 1. Detailed cloning procedures can be obtained from the authors.

### Cell culture

hTERT-RPE1 cell cultures were maintained in DMEM-F12 medium with 10 % fetal bovine serum (FBS), 5 U ml^−1^ penicillin and 50 μg ml^−1^ streptomycin at 37 and 5 % CO_2_

### Cell lines

All experiments were performed in hTERT-RPE1 cells. Stable RPE1 cellines were lentivirus generated pools. For all cell lines a PGK promoter was used. Third generation lentivirus was generated using procedures and plasmids previously described^40^. Briefly, the GFP or Cherry fusions were generated as Gateway ENTRY plasmids using standard molecular biology techniques. From these vectors, lentiviral transfer vectors were generated by recombination into Lentiviral “Destination” vectors derived from pCDH-EF1α-MCS-IRES-PURO (SystemBioSciences) using a Gateway LR reaction. VSV-G pseudotyped lentiviral particles were packaged using a third generation packaging system (Addgene plasmids number 12251, 12253 and 12259)^41^. Cells were then transduced with virus and stable expressing populations were generated by antibiotic selection. Used cell lines are listed in Supplemental Table 2.

### Immunostaining

Cells grown on coverslips were fixed with 3 % formaldehyde (Polyscience) for 15 minutes in room temperature and permeabilized with 0.05 % Saponin (Sigma-Aldrich) in PBS. Cells were then stained with the indicated primary antibodies for 1 hour, washed in PBS/Saponin, stained 1 hour with fluorescently labeled secondary antibodies and washed with PBS/Saponin. Cells were mounted using Mowiol (Sigma-Aldrich). Cells stained for Rab7 was pre-permeabilized in 0.05 % Saponin in PEM buffer (80 mM K-Pipes, 5 mM EGTA, 1 mM MgCl_2_ (pH 6.8)) for 5 minutes in room-temperature, before fixation in 3 % formaldehyde.

### Transient transfection

hTERT-RPE1 cells were transfected using Fugene 6 using 3 μl reagent per 1 μg of DNA. Cells for live cell imaging were transfected in MatTek dishes (Inter Instruments) dishes and cells for fixation were transfected in 24 wells with coverslips.

### siRNA

All siRNA used was obtained from Ambion. Cells were transfected using lipofetcamine RNAiMAX transfection reagent (Life Technologies) following the manufacturers protocol and 50 nM siRNA targeting WDFY2, (sense: GCAUGUCUUUUAACCCGGA) (s41881). Silencer^™^ select negative control No. 2 siRNA (AM4637) was used as a control.

### CRISPR/Cas9-mediated deletion of WDFY2

hTERT-RPE1 cells deleted for WDFY2 were generated using CRISPR/Cas9. Guide RNAs were designed using Benchling software (www.benchling.com). For deletion of WDFY2, a guide RNA binding in directly in front of the start codon in exon2 and one binding after exon2 was chosen (gRNA1: 5’-TGGATCTCCGCCGCCATCGG-3’; gRNA2: 5’-ACTACTGCCATTCGGCCGCG-3’). This strategy resulted in deletion of the first 46 amino acids of WDFY2 and deleted also the splice donor of the intron linking exon 2 and 3 (Figure S2). We reasoned that this deletion should not result in a functional cDNA due to deletion of the splice site.

pX458-derived plasmids encoding both Cas9-2A-GFP and the respective gRNA were transfected using Fugene6^20^. 48 hours post-transfection, GFP-positive cells were sorted and seeded out in several dilutions to obtain single colonies, which were picked and characterized. Clones lacking WDFY2 were identified by PCR, and the introduced mutations were characterized by PCR followed by cloning and sequencing.

## Microscopy

### Confocal fluorescence microscopy

Confocal images were obtained using LSM710 confocal microscope (Carl Zeiss) equipped with an Ar-laser multiline (458/488/514 nm), a DPSS-561 10 (561 nm), a laser diode 405-30 CW (405 nm), and a HeNe laser (633 nm). Images were taken using a Plan-Apochromat 63x/1.40 oil DIC III (Carl Zeiss).

### Live time-lapse microscopy

Live-cell imaging was performed on a Deltavision OMX V4 microscope equipped with three PCO.edge sCMOS cameras, a solid-state light source and a laser-based autofocus. Cells were imaged in Live Cell Imaging buffer (Invitrogen) supplemented with 20 mM glucose. Environmental control was provided by a heated stage and an objective heater (20-20 Technologies). Images were deconvolved using softWoRx software and processed in ImageJ/FIJI^42^.

### Structured illumination microscopy

RPE1 cells stabily expressing GFP-WDFY2 were transiently transfected with Cherry-Rab5/Rab4/Rab11, Cherry-VAMP3 or stained with anti Rab7/Vps26/Vps35, cells were fixed with 4 % PFA and 0.1 % glutaraldehyde stained with mouse-GFP (Roche) and RFP booster (Chromotek) or secondary antibodies then embedded in ProLongGold Antifade Reagent (Life Technologies). Three dimensonial SIM imaging was performed on Deltavision OMX V4 microscope with an Olympus x60 NA 1.43 objective and three PCO edge sCMOS cameras. Using 488 and 568 laser lines to excite the used fluorophores, cells were illuminated with a grid pattern and for each image plane, 15 images were acquired. Images were reconstructed from the raw image files, aligned and projected using Softworx software (Applied Precision, GE Healthcare). Images were processed in ImageJ/Fiji ^42^.

### STORM imaging

Cells stably expressing GFP-WDFY2 were fixed in 4 % PFA and labelled with anti-GFP. Imaging was performed in 100 mM Tris, 50 mM NaCl with an oxygen scavenging system (10 % Glucose, 10 kU catalase, 0.5 kU glucose oxidase) and 10 mM MEA as reducing agent. Imaging was performed using a Deltavision OMX V4 microscope (GE Healthcare); localization and reconstruction were performed in softWoRx software; all further image processing was performed in Fiji.

### DNA-PAINT microscopy

Cells were seeded in an 8 well chamber (Lab-Tek) and fixed with 4 % formaldehyde for 15 minutes in room temperature to preserve endosome tubules. Staining and imaging was done according to the manufacturers protocol (Ultivue). Imaging was performed using a Deltavision OMX V4 microscope (GE Healthcare); localization, reconstruction and alignment were performed in softWorRx software; all further image processing was performed in Fiji.

### Protein-Lipid overlay assays

In vitro lipid binding activities of full length WDFY2 were determined by protein-lipid overlay assay ^43^. Protein purification and lipid overlay assay was performed as previously described ^44^.

### GFP affinity purification

Stable hTERT-RPE1 cellines expressing GFP or GFP WDFY2 were seeded in 10 cm dishes up to 80 % confluence and then lysed in lysisbuffer containing, 50 mM TRIS, 150 m NaCl, 0,25 % Triton X100, 1 mM DTT, 50 μM ZnCl_2_, 5 mM NaPPi, 20 mM NaF, 1x of phosphatase inhibitor 3 (S/T), phosphatase inhibitor 2 (Y) and protease inhibitor mix. GFP-trap magnetic beads (ChromoTek) were added to the lysate and incubated rotating at 4 degrees for 4 hours.

### LC-MS/MS and Protein identification and label-free quantitation

Beads containing bound proteins were washed 3 times with PBS, reduced with 10 mM DTT for 1 hour at 56°C followed by alkylation with 30 mM iodoacetamide in final volume of 100 μl for 1 hour at room temperature. The samples were digested over night with Sequencing Grade Trypsin (Promega) at 37°C, using 1.8 μg trypsin. Reaction was quenched by adding 1 % trifluoracetic acid to the mixture. Peptides were cleaned for mass spectrometry by STAGE-TIP method using a C18 resin disk (3M Empore)^45^. All experiments were performed on a Dionex Ultimate 3000 nano-LC system (Sunnyvale CA, USA) connected to a quadrupole – Orbitrap (QExactive) mass spectrometer (ThermoElectron, Bremen, Germany) equipped with a nanoelectrospray ion source (Proxeon/Thermo). For liquid chromatography separation we used an Acclaim PepMap 100 column (C18, 2 μm beads, 100 Å, 75 μm inner diameter) (Dionex, Sunnyvale CA, USA) capillary of 25 cm bed length. The flow rate used was 0.3 μL/min, and the solvent gradient was 5 % B to 40 % B in 120 minutes, then 40-80 % B in 20 minutes. Solvent A was aqueous 2 % acetonitrile in 0.1 % formic acid, whereas solvent B was aqueous 90 % acetonitrile in 0.1 % formic acid.

The mass spectrometer was operated in the data-dependent mode to automatically switch between MS and MS/MS acquisition. Survey full scan MS spectra (from m/z 300 to 1,750) were acquired in the Orbitrap with resolution R = 70,000 at m/z 200 (after accumulation to a target of 1,000,000 ions in the quadruple). The method used allowed sequential isolation of the most intense multiply-charged ions, up to ten, depending on signal intensity, for fragmentation on the HCD cell using high-energy collision dissociation at a target value of 100,000 charges or maximum acquisition time of 100 ms. MS/MS scans were collected at 17,500 resolution at the Orbitrap cell. Target ions already selected for MS/MS were dynamically excluded for 45 seconds. General mass spectrometry conditions were: electrospray voltage, 2.0 kV; no sheath and auxiliary gas flow, heated capillary temperature of 250°C, heated column at 35°C, normalized HCD collision energy 25 %. Ion selection threshold was set to 1e5 counts. Isolation width of 3.0 Da was used.

MS raw files were submitted to MaxQuant software version 1.4.0.8 for protein identification^46^. Parameters were set as follow: protein N-acetylation, methionine oxidation and pyroglutamate conversion of Glu and Gln as variable modifications. First search error window of 20 ppm and mains search error of 6 ppm. Trypsin without proline restriction enzyme option was used, with two allowed miscleavages. Minimal unique peptides were set to 1, and FDR allowed was 0.01 (1%) for peptide and protein identification. Label-free quantitation was set with a retention time alignment window of 3 min. The Uniprot human database was used (downloaded august 2013). Generation of reversed sequences was selected to assign FDR rates.

### GFP pulldown assay

GFP trap (GFP-trap magnetic beads, ChromoTek) was used for interaction studies and the experiments were performed according to the manufacturer’s protocol. Stable RPE1 cell lines expressing GFP–VAMP3 in combination with Cherry-WDFY2 were used. Stable RPE1 cells expressing Cherry-WDFY2 transiently transfected with pEGFP-N2 were used as control. Five per cent input was used for the immunoblotting of the GFP traps.

### VAMP3 redistribution assay

Cells were transiently transfected with GFP-VAMP3 and Cherry-Cortactin overnight and fixed in 3 % PFA. Cells were labelled with anti-GFP (Roche) and RFP-booster (Chromotek) Images were acquired on LSM710 Confocal microscope using a Plan-Apochromat 63x/1.40 oil DIC III (Carl Zeiss) objective. Cells were selected in the red channel (cortactin). Images were processed in Fiji using a custom Python script.

### Measuring compartment-specific VAMP3 localization

RPE1 (WT) and WDFY2 (-/-) cells were transfected with GFP-Vamp3 and fixed in 3 % PFA. Cells were labelled with human anti-EEA antibody and rabbit anti-Lamp1 antibody and corresponding fluorophore-conjugated secondary antibodies. Images were acquired using a LSM710 confocal microscope using a Plan-Apochromat 63x/1.40 oil DIC III (Carl Zeiss) objective. To measure VAMP3 distribution, EEA1 and Lamp1 vesicles were automatically segmented and GFP-Vamp3 intensity was measured within the respective compartments using a custom Python-based Fiji script^42^. Individual measurements were then collected and plotted.

### Recycling and exocytosis experiments

RPE1 (WT) and WDFY2 (-/-) cells were seeded in MatTek dishes (Inter Instruments) and transfected with pHluorin-VAMP3 or pHluorin-MT1-MMP. Imaging was performed on Deltavision OMX V4 microscope (GE Healthcare) using a 60x TIRF objective. Images were taken every second for two minutes. Images were manually processed using Fiji by scoring bright dots that appeared and disappeared within a few frames^42^.

### Gelatin degradation assay

Oregon Green-conjugated gelatin-coated (Life Technologies) coverslips were prepared as described previously with some modifications^25^. Briefly, coverslips (12 mm diameter, No. 1 thickness, VWR international) were pre-cleaned in 20 % nitric acid overnight. After extensive washing, the coverslips were coated with 50 μg/mL poly-l-lysine (Sigma-Aldrich) for 30 minutes, washed in PBS, and fixed with cold 0.5 % glutaraldehyde (Sigma-Aldrich) in PBS for 15 minutes on ice. Subsequently, the coverslips were washed in PBS and coated for 20 minutes with prewarmed 10 mg/mL Oregon Green-conjugated gelatin / 2 % sucrose in PBS. After coating, the coverslips were washed with PBS and incubated in 5 mg/mL sodium borohydride (Sigma-Aldrich) for 15 minutes. The coverslips were then washed with PBS, sterilized with 70 % ethanol, and equilibrated in serum-containing medium for 1 hour before the addition of cells. For Gelatin degradation assays, 5×10^4^ RPE1 (WT) and WDFY2 (-/-) cells were suspended in 1 ml culture medium and added to wells with gelatin coated cover slips followed by 6 hours incubation at 37°C. The cells were then fixed in 3 % formaldehyde in PBS for 15 minutes, permeabilized with 0.1 % Triton X-100 (Sigma-Aldrich) in PBS, incubated with Rhodamine phalloidin (Life Technologies) for 15 minutes and mounted for examination by confocal microscopy. Cells incubated with MMP inhibitor GM6001 (VWR international, J65687.MX) were seeded in growth medium containing the MMP inhibitor.

Samples were investigated using a LSM710 confocal microscope (Carl Zeiss), a 63x objective and zoom 1.0. Cells / field of imaging were chosen on basis of the nuclear staining and gelatin quality, and at least 15 images were randomly taken throughout the cover slips in each experiment, giving at least 120 cells per experiment for each condition. All images within one experiment were taken with constant gain and pin-hole parameters.

### Inverted Matrigel^™^ invasion assay

Inverted invasion assays were performed as previously described^47^. In brief, Matrigel^™^ (Corning) was supplemented with 25 μg/ml fibronectin (Sigma-Aldrich) and 80 μl was added to Transwell^®^ (Sigma-Aldrich, 8 μm pores) and allowed to polymerize for 45 minutes at 37°C. The inserts were then inverted and 4×10^4^ cells were seeded on top of the filter on the opposite side from the Matrigel. The transwells were placed in serum free medium and the upper chamber was filled with serum supplemented medium (10 % v/v FBS) with HGF (100 ng/ml). 72 hours after seeding (48 hours after seeding in the case of knockdown) cells were stained with Calcein AM (4 μM) (Thermo Fisher) for 1 hour before invading cells were visualized with confocal microscopy (Zeiss LSM 710, ×20 objective). Cells that did not make it through the filter were removed with a tissue paper. Section of 10 μm intervals (for quantification) and 1.23 μm intervals (for 3D reconstruction) were captured. Images were analyzed with Fiji^42^. Invasion is presented as the sum of white pixels of all slides from 50 μm and beyond, divided by the sum of white pixels of all slides. a 3D reconstruction of z stacks was done in Paraview (https://www.paraview.org/).

### Actin depolymerization assay

Cells stably expressing GFP-WDFY2 were seeded in MatTek dishes (Inter Instruments). Cells were imaged like described for Live time-lapse microscopy. Images were acquired every 3 seconds. Cells were imaged for a period before Latrunculin B (Merck, 428020) was added with a final concentration of 10 μM.

### Quantitative real-time PCR of mRNA expression

mRNA expression analysis was done as described in Pedersen et al, 2012^48^. The primers used in the experiment were QT00035455 for WDFY2 and QT00000721 for TBP for housekeeping (Qiagen).

### Image processing and data analysis

All live cell images were deconvolved using Softwork (GE Healthcare) prior to analysis and presentation. All further image analysis and measurements steps were performed in Fiji using custom Python scripts^42^. Postprocessing of data was performed in Python using the Pandas and Seaborn packages^49^. All used scripts, as well as all raw data required to reproduce the plots shown in the manuscript are available on GitHub (https://github.com/koschink/Sneeggen_et_al).

### Statistics

Statistical analysis was performed using Graphpad Prism and the Scipy “Stats” package. Student’s t-test was used as a measure for statistical significance in samples with a gaussian distribution. For analysis of multiple samples, we utilized ANOVA and groups were compared using Bonferroni’s Multiple Comparison Test. ^∗^P < 0.05, ^∗∗^P < 0.01, ^∗∗∗^P < 0.001.

## Acknowledgements

We are grateful to Philippe Chavrier, Matthew Seaman and Jim Norman for kindly providing plasmid constructs. We thank Anne Engen for help with cell culture, Chema Bassols for IT assistance, and Eva Rønning and Pepijn Wopken for laboratory assistance. The core facilities for Advanced Light Microscopy and Proteomics at Oslo University Hospital are acknowledged for assistance with microscopes and proteomic analyses, respectively. We thank Trine Håve, Ulrikke Dahl Brinch, Kia Wee Tan and Hélène Spangenberg for assistance with plasmid constructs and characterization of CRISPR clones and Ling Wang for preparing gelatin coated coverslips. K.O.S. and E.M.H. hold career development fellowships and N.M.P a postdoctoral fellowship from the Norwegian Cancer Society. H.S. is supported by a grant from the South-Eastern Norway Regional Health Authority. C.C. holds a Young Research Talents grant from the Research Council of Norway. This work was partly supported by the Research Council of Norway through its Centres of Excellence funding scheme, project number 262652.

## Author Contributions

K.O.S and H.S supervised the study. M.S and K.O.S conceived the study and designed experiments. M.S generated construct, lentivirus and stable cell lines, performed super resolution imaging and all live cell imaging, GFP trap experiment, VAMP3 distribution experiment, the compartment specific Vamp3 distribution experiment, exocytosis experiment, invasion experiments, image analyzing and quantifications. K.O.S wrote image and data processing software and analysed data, performed CRISPR/Cas9 knockouts, generated construct, lentivirus, stable cell lines and helped with STORM imaging. N.M.P performed GFP-pulldown experiments confirming interactions and designed, performed and analyzed the gelatin degradation assays. C.C performed GFP-trap experiment and analyzed mass spectrometry data. E.M.H helped with the invasion assay. M.S, K.O.S. and H.S wrote the manuscript with input from all co-authors.

---

**Supplemental Figure 1:**
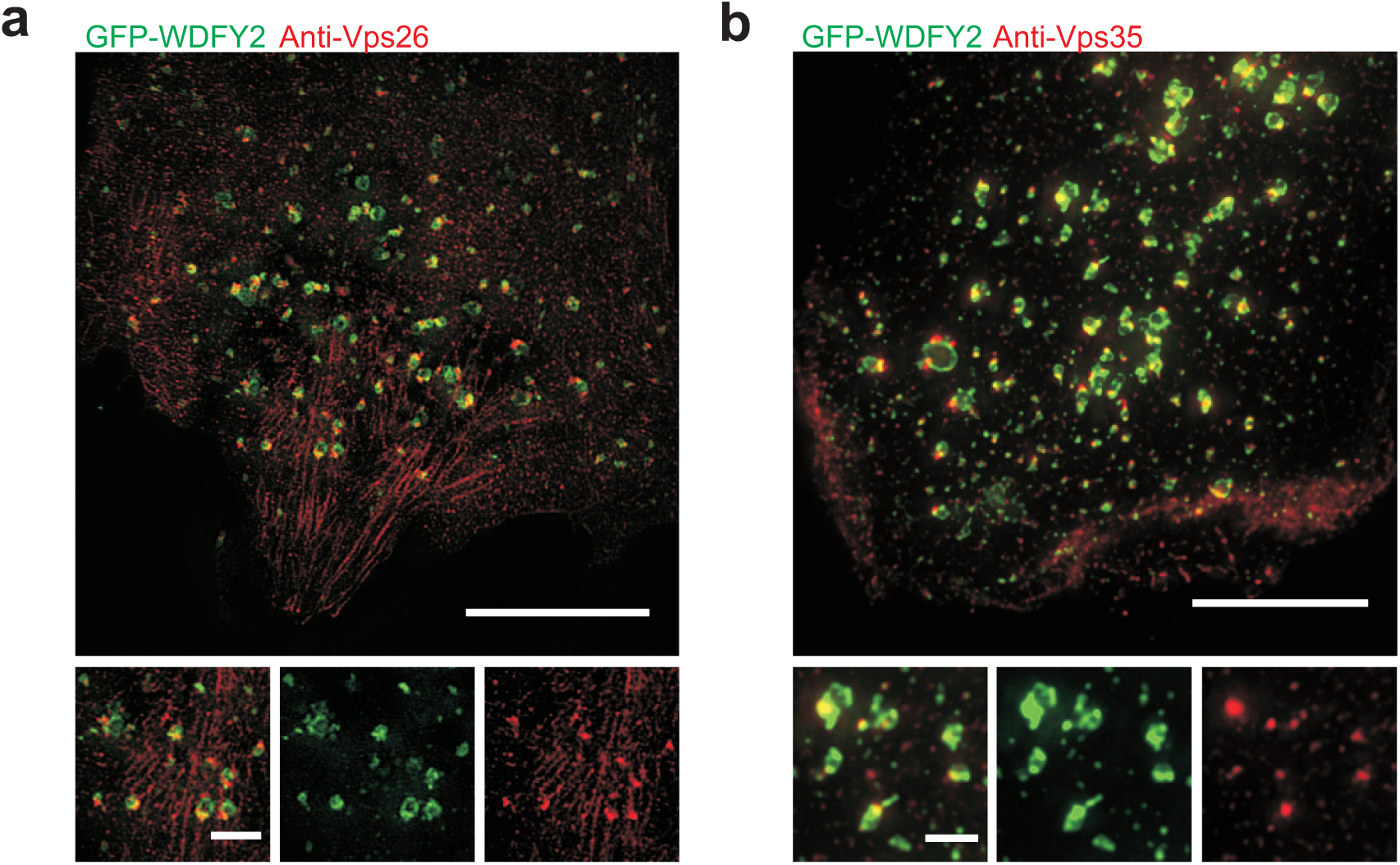
a. SIM image showing GFP-WDFY2 and anti-Vps26 visualized with antibody localization to endosomes and the base of WDFY2 positive tubules. Scale bar overview image: 10 μm, scale bar of inset: 1 μm (n=8 cells).
b. SIM image showing GFP-WDFY2 and Vps35 visualized with antibody localization to endosomes and the base of WDFY2 positive tubules. Scale bar overview image: 10 μm, scale bar of inset: 1 μm (n=8 cells).

**Supplemental Figure 2:**
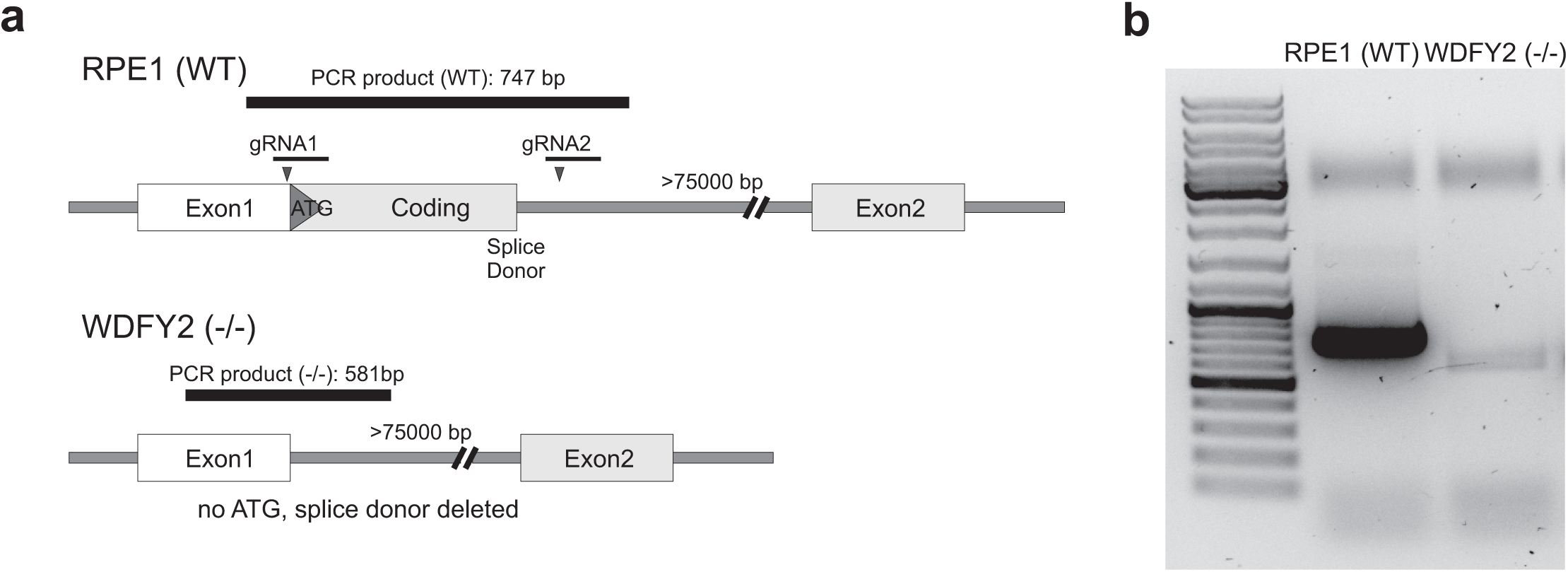
a. Schematic presentation of the CRISPR/Cas9 -generated WDFY2 deletion. Indicated are location of the guide RNAs and the primers used to characterize the resulting clones. The knockout was designed that the coding region including the starting ATG and the splice donor of exon 2 was completely excised, preventing the generation of a functional mRNA.
b. PCR characterization of the CRISPR/Cas9 -generated WDFY2 deletion. Expected band sizes: RPE (WT): 747 bp, WDFY2 (-/-): 581 bp.

**Supplemental Figure 3:**
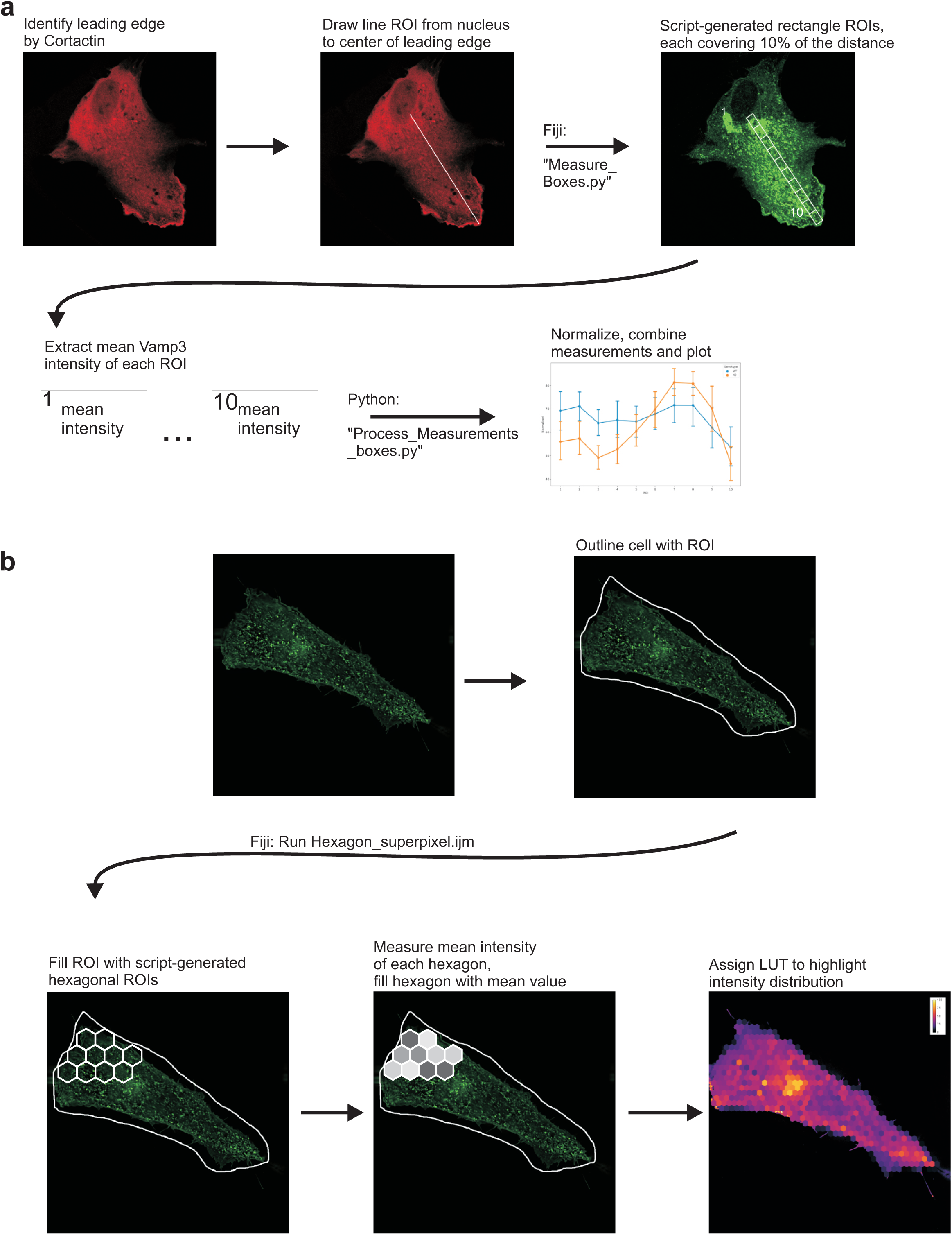
a. Schematic display of the processing steps to extract the data displayed in Figure 4b. Cells expressing GFP-Vamp3 were stained with Cortactin to identify leading edges. A line ROI was drawn between the rim of the nucleus and the leading edge, and rectangular ROIs, each covering 10% of the distance between the leading edge, were automatically generated using the Imagej script “Measure_Boxes.py” and their mean intensity in the Vamp3 channel was extracted. Mean intensities of all cells were normalized and plotted using the script “Process_Measurements_boxes.py” in Python.
b. Generation of superpixels to visualize Vamp3 distribution. Cells were outlined using ImageJ and an ImageJ macro (https://gist.github.com/mutterer/035ade419bf9c96475ce) was used to generate hexagonal ROIs. Mean intensity values for each hexagon were measured and the corresponding hexagon filled with the mean value, thereby generating superpixels. A false-color lookup table was applied to visualize intensities.

**Supplemental Figure 4:**
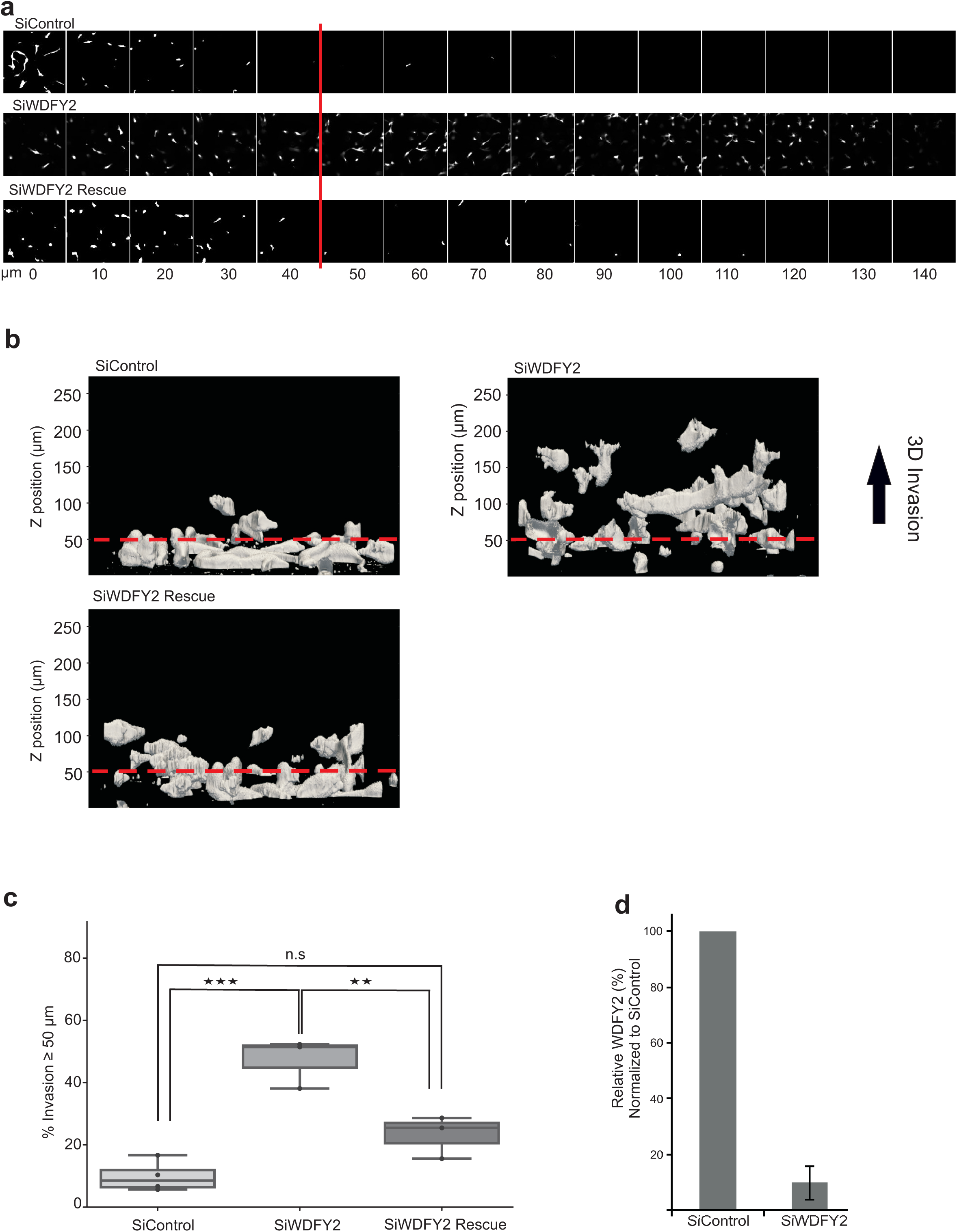
a. Optical sections (10 μm) of RPE1 cells transfected with non-targeting siRNA (SiControl) and siRNA targeting WDFY2 (SiWDFY2) and RPE1 cells expressing siRNA resistant GFP-WDFY2 transfected with siRNA targeting WDFY2 (siWDFY2 rescue) invading fibronectin-supplemented Matrigel^™^. Cells were stained with Calcein-AM. The red line indicates the z-axis threshold (50 μm) defining invading cells.
b. Orthogonal view of a 3D reconstruction of RPE1 and RPE1 (WDFY2 rescue) cells treated as above. The red line indicates the z-axis threshold (50 μm) defining invading cells.
c. Quantification of invasion of RPE1 cells treated as above. Invasion was quantified as described in the experimental section. Plotted data points indicate measurements derived from individual confocal z-stacks, color-coded per experiment (15 z-stacks per experiment, n= 3/4 experiments). ANOVA with Bonferroni’s Multiple Comparison Test, p=0.0004.
d. Knockdown efficiency of WDFY2 is shown by quantitative realtime PCR. Shown as relative to control. Error bars denote +/-S.D (n= 4)

**Movie S1:** Deconvolved widefield time-lapse showing endosome labelled with GFP-WDFY2 and endosomal tubules positive for WDFY2. Images were acquired every 2 seconds. Scale bar: 1 μm.

**Movie S2:** Deconvolved widefield time-lapse showing endosome labelled with GFP-WDFY2 and mCherry-Coronin. Images were acquired every 2 seconds. Scale bar: 1 μm.

**Movie S3:** Deconvolved widefield time-lapse showing endosomes and endosomal tubules labelled with GFP-WDFY2 prior to addition of Latrunculin and after addition. Images were acquired every 3 seconds. Scale bar:10 μm.

**Movie S6:** Deconvolved widefield time-lapse showing endosome labelled with GFP-MT1-MMP and mCherry-WDFY2. Images were acquired every 2 seconds. Scale bar: 1 μm.

**Movie S5:** TIRF time-lapse movie showing secretion events of pHluorin-Vamp3 in RPE (WT) and WDFY2 (-/-). Images were acquired every second. Scale bar: 10 μm, scale bar of inset: 1 μm.

**Movie S6:** TIRF time-lapse movie showing exocytosis events of pHluorin-MT1-MMP in RPE (WT) and WDFY2 (-/-). Images were acquired every second. Scale bar: 10 μm, scale bar of inset: 1 μm.

